# Effective connectivity between the medial temporal lobes and early visual cortex modulated by unrestricted viewing predicts memory retrieval and gaze reinstatement

**DOI:** 10.1101/2025.02.18.638908

**Authors:** N. Ladyka-Wojcik, Z.-X. Liu, J.D. Ryan

## Abstract

Memory and gaze behavior are intricately linked, guiding one another to extract information and create mental representations of our environment for subsequent retrieval. Recent findings from functional neuroimaging and computational modeling suggest that reciprocal interactions between the extended hippocampal system and visuo-oculomotor regions are functionally relevant for building these mental representations during visual exploration. Yet, evidence for the directionality of information flow during encoding within this reciprocal architecture in humans is limited. In the current study, we used dynamic causal modeling (DCM) to give a non-invasive account for the directional influences between these systems when new memories are created. Here, we provide novel evidence demonstrating how unrestricted, naturalistic visual exploration induces changes in this connectivity. Subsequent memory retrieval performance was also predicted by the pattern of connectivity modulated by unrestricted visual exploration, identifying the mechanism underlying a rich history of previous work linking increased gaze behavior during encoding to later memory. Together, these findings suggest that gaze behavior shapes the ways in which brain dynamics within and between the hippocampal system and early visual cortex unfold during encoding in humans. Importantly, these directional interactions support the building of coherent, lasting mental representations.

## 1 Introduction

What we encode in memory is intimately tied to where we look (Hannula et al., 2010; Ryan & Shen, 2020). Visual exploration allows us to accumulate salient information from our environment to build mental representations in support of memory retrieval (Liu et al., 2017; Loftus, 1972; Nikolaev et al., 2023; Sharot et al., 2008). During encoding, a greater number of gaze fixations—the discrete stops made by the eyes—predicts mnemonic performance (Damiano & Walther, 2019; Loftus, 1972). Conversely, when fixations are explicitly restricted, subsequent memory retrieval is impaired (Damiano & Walther, 2019; Henderson et al., 2005; Liu et al., 2020). Patterns of gaze fixations at encoding (i.e., scanpaths) are also spontaneously reproduced during retrieval in support of memory (Noton & Stark, 1971a, 1971b; for a review, see Wynn et al., 2019). Notably, where we move our eyes during encoding is not solely driven by bottom-up, exogenous features of space; visual exploration is also guided by top-down factors such as prior knowledge and memories of past experiences (Castelhano & Henderson, 2007; Ramey et al., 2020; Ryan et al., 2007). Eye movements increasingly anticipate upcoming stimulus locations over the course of learning which, paired with a shift in brain activity from representing previously viewed images to upcoming ones, predict later successful memory recall (Büchel et al., 2024; Huang et al., 2023). Thus, the neural architecture of the human brain must support the rapid exchange of information about mental representations and visual exploration, in service of cognition and behavior.

The hippocampal (HPC) memory and visuo-oculomotor control systems have been widely investigated for their respective roles in memory (Eichenbaum et al., 1994, 1996; Moscovitch et al., 2016) and gaze behavior (Robinson, 1986; Robinson & Optican, 1981). Recently, vast structural and functional connections between these systems have been identified in the human and nonhuman primate brain (e.g., Shen et al., 2016). Whereas early theoretical accounts positioned the HPC as the site at which already-parsed sensory information is bound into lasting representations in memory (Cohen & Eichenbaum, 1993), more recent accounts propose that connections between these systems might allow for eye movements themselves to be embedded into mnemonic representations (Katz et al., 2020; Ryan, 2024). Compelling evidence for this notion comes from computational modeling, which has revealed numerous polysynaptic projections between these systems along which information may flow (Shen et al., 2016). Indeed, simulating stimulation to HPC subfields and the broader medial temporal lobes (MTL) resulted in propagating signals via multiple regions, including visual regions, that ultimately reached oculomotor control regions within the period of a gaze fixation (Ryan et al., 2020). Converging findings from electrophysiological recordings have identified neuronal responses in the HPC tuned to visual exploration in nonhuman primates (Hoffman et al., 2013; Jutras et al., 2013; Mao et al., 2021) and in humans (Katz et al., 2022). Moreover, amnesic cases with damage to the HPC system show selective deficits in their gaze patterns (Ryan et al., 2000; Warren et al., 2011) such as less organized viewing patterns relative to healthy individuals (Lucas et al., 2019). Together these findings speak to the existence of reciprocal connections between the HPC memory and visuo-oculomotor control systems, which could coordinate encoding-relevant viewing behavior towards previously seen and new locations (Knapen, 2021; Nikolaev et al., 2024; Ryan & Shen, 2020).

A question remains: how are the strength and directionality of connections between the HPC memory and visuo-oculomotor systems modified by viewing behavior during encoding? To this end, measures of effective connectivity offer the ability to model *directed influences* between these systems due to modulatory effects (Friston et al., 2003). Previous work by Tullo et al. (2023) found that perception of highly familiar scenes evokes bottom-up excitatory and top-down inhibitory effective connectivity between visual regions and subregions of the parahippocampal place area (PPA), a key area of the extended HPC memory system implicated in scene processing (Aguirre et al., 1996; Epstein et al., 1999; Epstein & Baker, 2019; Epstein & Kanwisher, 1998). This finding highlights plausible pathways between the HPC memory and visuo-oculomotor systems, but it is unknown if we should expect similar patterns of effective connectivity during encoding when new visuo-spatial information must be bound into mnemonic representations. Moreover, without manipulating visual behavior, we cannot discern whether gaze fixations directly modulate the strength and directionality of these pathways. Multi-modal approaches leveraging functional neuroimaging (i.e., fMRI) and simultaneous eye-tracking have provided arguably the strongest insights into this question to date. Functional connectivity between the HPC, PPA, and early visual regions during the encoding of novel scenes was impeded when fixations were actively restricted compared to unrestricted, free, viewing (Liu et al., 2020). These effects were stronger for real-world scenes compared to scrambled color-tile images, for which there was relatively less information available for extraction, and the effects were also stronger for scenes that were subsequently remembered versus forgotten. A recent effective connectivity study supplies evidence consistent with this latter finding: successful encoding of novel scenes was associated with excitatory effective connectivity from the PPA to the HPC (Schott et al., 2023), although regions of the visuo-oculomotor system were not considered. At the level of the visuo-oculomotor system, functional activation in the occipital pole (OCP) has been tied to the formation and later reinstatement of gaze fixations made during *free*-viewing of novel scenes (Wynn et al., 2022). The authors also identified voxels in the HPC and PPA responsive to both subsequent memory strength and gaze reinstatement, suggesting that information about eye movements may be embedded into mnemonic representations. However, it is unknown whether effective engagement of the extended HPC and visuo-oculomotor systems differs based on subsequent memory and/or gaze reinstatement. In fact, to our knowledge, viewing behavior during encoding has not yet been considered in studies of effective connectivity between these systems.

In the present study, we used the data from Liu et al. (2020) to conduct Dynamic Causal Modeling (DCM), in order to probe the top-down and bottom-up modulatory nature of visual exploration during novel scene encoding in the healthy adult brain. Specifically, our main question of interest focused on how unrestricted, naturalistic visual exploration (i.e., *free*-viewing) during scene and scrambled image encoding could modulate the effective connectivity architecture between the HPC, PPA, and OCP, over-and-above restricted (i.e., *fixed*-) viewing. Briefly, DCM models the (1) underlying (i.e., average) effective connectivity within- and between-regions of interest (ROIs), (2) experimental modulation of within-(i.e, self-), forward, and backward connections of ROIs, and (3) the external input that stimulates the system. Here, we compared families of models with top-down, bottom-up, and bidirectional connections switched on or off, evaluating underlying and modulatory effects of *free*-viewing for scene and scrambled image encoding. We predicted that the modulatory effects of unrestricted gaze fixations during encoding would be evidenced in strengthened excitatory connections from the early visual cortex towards the PPA for both scenes and scrambled images. This hypothesized architecture would facilitate bottom-up information flow accumulated through visual exploration, from the visuo-oculomotor system to the extended HPC memory system. During unrestricted viewing of scene images, when meaningful visuo-spatial information can be extracted to build mental representations for later retrieval, we hypothesized that this excitatory effect would propagate from the PPA towards the HPC. We also hypothesized a top-down, inhibitory modulatory effect of unrestricted viewing from the HPC to the PPA, thought to reflect suppression of mental representations from previously encoded images. Finally, we interrogated whether there were specific modulatory effects that predicted memory strength and/or similarity of gaze reinstatement during subsequent retrieval of scenes encoded under unrestricted viewing conditions. Importantly, we considered both self- and between-region modulatory effects, as both are integral to understanding the excitatory and inhibitory balance in the modeled architecture (see for e.g., Snyder et al., 2021). Our findings demonstrate how visual exploration dynamically modifies the connections between the extended HPC memory and visuo-oculomotor systems to support the rapid formation of scene representations in memory.

## 2 Materials and Methods

### 2.1 Participants

Our participants consisted of thirty-six young adults (22 females) aged 18-35 years (*M* = 23.58 years, *SD* = 4.17) from the University of Toronto and surrounding Toronto area (education: *M* = 16.27 years, *SD* = 1.8) reported in Liu et al. (2020). Participants had normal or corrected-to-normal vision (including color vision) and were screened for any neurological or psychological conditions. The study was approved by the Research Ethics Board at the Rotman Research Institute at Baycrest Health Sciences, and all participants provided written informed consent.

### 2.2 Stimuli

A comprehensive description of the stimuli used in this experiment is reported in Liu et al. (2020). Briefly, stimuli consisted of twenty-four color scene images (500 x 500 pixels; viewing angle: 7.95 x 7.95 degree) for each of 36 semantic scene categories (e.g., warehouse, movie theater, airport, etc.), for a total of 864 images. Images were either obtained from a previous study by Park et al. (2015) or from Google Images using the same semantic categories as the search term. Images in each semantic scene category (8 images/category) were randomly assigned to either the *free*-viewing encoding condition, the *fixed*-viewing encoding condition, or a retrieval task condition not reported here (see Liu et al., 2020). The assignment of the images to these experimental conditions was counterbalanced across participants, with low-level features such as luminance equalized across conditions in MATLAB using the SHINE toolbox (Willenbockel et al., 2010).

Next, the eight images for each scene category in the *free*- and *fixed*-viewing conditions were randomly assigned to eight fMRI encoding runs, such that each run contained a scene from each category shown under *free*- and *fixed*-viewing conditions. Finally, images from two randomly selected fMRI runs were scrambled, using six levels of tile sizes (i.e., 5, 10, 25, 50, 100, and 125 pixels) to resemble the clutter feature of the scene images. The pixels within each tile were averaged to create two runs of scrambled color-tile images.

Stimuli were presented with Experiment Builder (Eyelink 1000; SR Research) back projected to a screen (projector resolution: 1024×768) and viewed with a mirror mounted on the MRI head coil.

### 2.3 Procedure

Participants completed eight runs of the encoding task, with six runs containing scene images and two runs containing scrambled images (Figure 1). Each run had 72 images, studied under *free-*viewing instructions (i.e., 36 trials) or *fixed-*viewing instructions (i.e., 36 trials). The order of the scene and scrambled image runs was randomized across participants. Within an encoding trial, participants first saw a fixation cross (‘+’) for 1.72-4.16 secs whose location was randomly determined within a radius of 100 pixels around the center of the screen. The fixation cross was presented in either a green color (indicating the subsequent trial image was in the *free-*viewing condition) or in a red color (indicating the subsequent trial image was in the *fixed-*viewing condition). For the *free-* viewing condition, participants were instructed to freely explore the scene or scrambled image that followed with their eyes in order to encode as much information as possible. For the *fixed-*viewing condition, participants were instead instructed to keep their eye gaze at the location of the fixation cross for the duration of the presented scene or scrambled image that followed. Each run was 500 seconds long, with 10 seconds and 12.4 seconds added to the start and end of the task, respectively. The trials from the two viewing conditions (i.e., *free*- and *fixed*-viewing) were pseudo-randomized to obtain an adequate design efficiency by choosing the design with the best efficiency from 1000 randomizations using MATLAB code by Spunt (2019). Please see Liu et al. (2020) for further details about the encoding task procedure.

**Figure 1.**
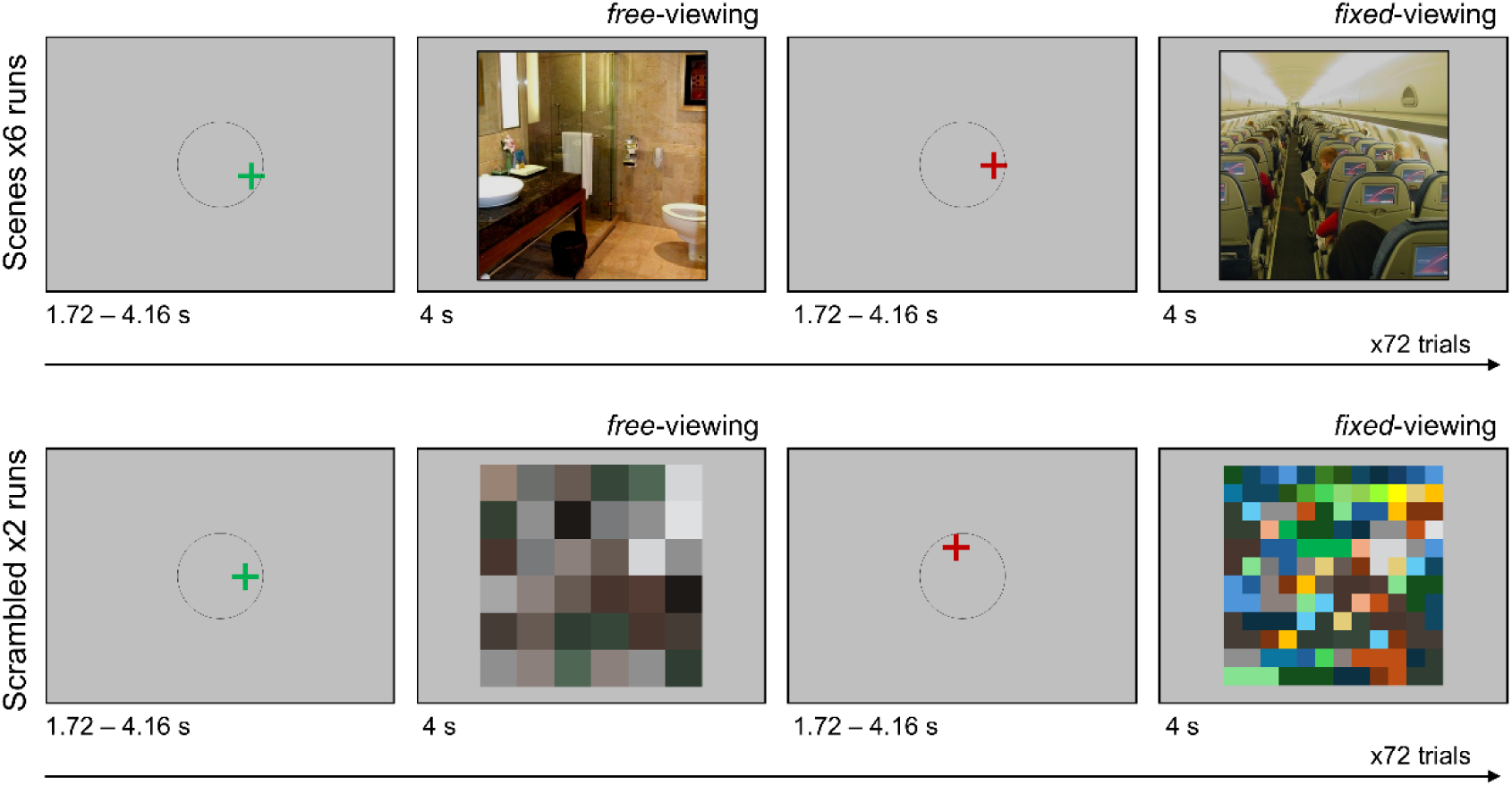
Visualization of experimental task procedure, adapted from Liu et al. (2020). Participants were presented with either a green or red fixation cross within a radius of 100 pixels around the center of the screen at the start of each encoding trial. A green fixation cross instructed participants to freely view the upcoming scene or scrambled color-tile image (*free*-viewing condition). A red fixation cross instructed participants to maintain fixation on the location of the cross during the presentation of the upcoming scene or scrambled color-tile image (*fixed*-viewing condition). Participants were presented with 6 runs of scenes and 2 runs of scrambled color-tile images, the order of which was randomized for each participant.

### 2.4 Eye-tracking

During the encoding task, monocular eye movements (right pupil and corneal reflection) were recorded inside the scanner using the EyeLink 1000 MRI-compatible remote eye-tracker with a 1000 Hz sampling rate (SR Research LTD., Ottawa, Ontario, Canada). At the start of the experiment, calibration was performed using the built-in EyeLink 9-point calibration procedure, with drift correction used between trials when necessary to ensure good tracking accuracy.

EyeLink’s default eye movement event parser was used to categorize fixations and saccades. Saccades were classified based on a velocity threshold of 30°/s and an acceleration threshold of 8000°/s (saccade onset threshold = 0.15°). Events not defined as saccades or blinks were classified as fixations. For each participant, the number of fixations made during the encoding task for each image was calculated and exported using EyeLink software Data Viewer.

### 2.5 MRI Data Acquisition

As specified in Liu et al. (2020), a 3T Siemens MRI scanner with a standard 32-channel head coil was used to acquire structural and functional MRI images. T1-weighted high-resolution MRI images for structural scans were obtained using a standard 3D MPRAGE (magnetization-prepared rapid acquisition gradient echo) pulse sequence (176 slices, FOV = 256 × 256 mm, 256 × 256 matrix, 1 mm isotropic resolution, TE/TR = 2.22/2000 ms, flip angle = 9°, and scan time = 280 s). For functional scans, BOLD signal was assessed using a T2*-weighted EPI acquisition protocol with TR = 2000 ms, TE = 27 ms, flip angle = 70 degrees, and FOV = 191 x 192 ms with 64 x 64 matrix (3 mm x 3 mm in-place resolution; slice thickness = 3.5 mm with no gap). A total of 250 volumes were acquired for the fMRI run, with the first 5 discarded to allow the magnetization to stabilize to a steady state. Both structural and functional images were acquired in an oblique orientation 30° clockwise to the anterior–posterior commissure axis.

### 2.6 fMRI Data Processing

The fMRI preprocessing procedure was previously reported in Liu et al. (2020); for completeness, it is re-presented here. Functional images were processed using SPM12 (Statistical Parametric Mapping, Wellcome Trust Center for Neuroimaging, University College London, UK; https://www.fil.ion.ucl.ac.uk/spm/software/spm12/; Version: 7771) in MATLAB (The MathWorks, Natick, USA), starting with the standard SPM12 preprocessing procedure. To specify, slice timing was first corrected using sinc interpolation with the reference slice set to the midpoint slice. Next, functional images were aligned using a linear transformation, and for each participant functional image parameters from the alignment procedure (along with global signal intensity) were checked manually using the toolbox ART (http://www.nitrc.org/projects/artifact_detect/). Anatomical images were co-registered to the aligned functional image and segmented into white matter, gray matter, cerebrospinal fluid, skull, and soft tissues using SPM12’s default 6-tissue probability maps. Segmented images were then used to calculate the transformation parameters mapping from the subjects’ native space to the MNI template space. The resulting transformation parameters were used to transform all functional and structural images to the MNI template. The functional images were finally resampled at 2 × 2 × 2 mm resolution and smoothed using a Gaussian kernel with an FWHM of 6 mm. The first five fMRI volumes from each run were discarded to allow the magnetization to stabilize to a steady state.

We then used SPM12 to conduct the first (i.e., subject) level whole brain voxel-wise General Linear Model (GLM) analysis for subsequent time series extraction. We included 4 conditions: all scene images, *free*-viewing scenes, all scrambled images, and *free*-viewing scrambled images, separately convolved trials with duration = 4 s with the canonical hemodynamic function (HRF). Motion parameters (6 from SPM realignment, 1 from ART processing), as well as outlier volumes, were added as regressors of no interest. Default high-pass filter with cut-off of 128 secs was applied. The serial correlation in fMRI time series in the restricted maximum-likelihood estimation of the GLM was accounted for using a first-order autoregressive model AR(1).

Since we were interested in determining how effective connectivity changes as a result of *free*-viewing over and above encoding under any viewing conditions, we constructed a *t* contrast for all scene trials and a *t* contrast for only *free*-viewing trials of scenes, averaged across all 6 scene runs. We did the same for the two runs of scrambled trials. Finally, we added an effect of interest *F*-contrast with the 4 conditions to extract time series for subsequent analyses of effective connectivity.

### 2.7 Time series extraction

Selecting ROIs for DCM analyses which show univariate contrast effects between conditions of interest is strongly recommended, in order to ensure that the ROIs under investigation are likely to play a role in the cognitive process targeted by the independent variable manipulation (Stephan et al., 2010; Zeidman et al., 2019a). As such, we used the significant group-level activation clusters to extract time series for the DCM analysis, which were constrained to be within the boundaries of our ROIs.

Specifically, we focused on three *a priori* ROIs – the HPC, PPA, and OCP – based on their involvement in scene encoding, visual exploration, and gaze reinstatement as reported in Liu et al. (2020) and Wynn et al. (2022). Subject-specific anatomical masks were generated for the HPC using FreeSurfer’s recon-all function (https://surfer.nmr.mgh.harvard.edu/; Fischl, 2012). The PPA was defined functionally by contrasting the scene and scrambled image conditions, collapsing over the *free*- and *fixed*-viewing conditions (Liu et al., 2020). For the OCP, we used peak activation coordinates reported in Wynn et al. (2022) to identify the center of an 8-mm radius spherical mask for each participant. Due to the increased complexity in interpretation when including inter-hemispheric connections for DCM, in part because testing the full model space would require additional nested models (Stephan et al., 2010; Stephan & Friston, 2010; Zeidman et al., 2019b), we combined left and right hemisphere masks.

Using these bilateral masks, we separately extracted voxels for the scene and scrambled image conditions that showed an effect of *free*-viewing (*p* = .05, no corrections). Importantly, including voxels that show task modulation effects can facilitate DCM analysis (Zeidman et al., 2019a). Therefore, when the number of voxels was lower than 50 (i.e., too few voxels in the ROI) for a specific ROI of a participant, we relaxed the threshold to 0.1, 0.5, or without using any threshold (i.e., threshold = 1) until at least 50 voxels could be obtained for that ROI and that participant (see Table 1). The average number of voxels (1 × 1 × 1 mm^3^) for the PPA, HPC, and OCP are reported in Table 1 below for both scene and scrambled image conditions.

**Table 1.** Averages and standard error of the mean (SE) for the number of voxels included in each condition and ROI, with voxel threshold counts across participants.

| ROIs | Scene |  |  |  |  | Scrambled |  |  |  |  |
| --- | --- | --- | --- | --- | --- | --- | --- | --- | --- | --- |
| | $M_{\text{voxels}}$ (SE) | Thresholds | | | | $M_{\text{voxels}}$ (SE) | Thresholds | | | |
|  |  | 0.05 | 0.1 | 0.5 | 1 |  | 0.05 | 0.1 | 0.5 | 1 |
| OCP | 252.72<br>(24.53) | 21 | 2 | 8 | 5 | 245.03<br>(26.32) | 17 | 4 | 5 | 10 |
| PPA | 477.78<br>(28.54) | 30 | 1 | 5 | 0 | 276.03<br>(31.14) | 20 | 4 | 12 | 0 |
| HPC | 354.25<br>(55.05) | 28 | 3 | 5 | 0 | 228.19<br>(32.51) | 22 | 9 | 5 | 0 |

### 2.8 Dynamic causal modeling

DCM is a framework for specifying models of effective connectivity among brain regions, estimating their parameters and testing hypotheses (Zeidman et al., 2019a). Since DCM estimates region-specific hemodynamic responses and neural states (Stephan & Friston, 2010), it is relatively resistant to issues related to HRF variations in different brain regions (Friston et al., 2014; Stephan et al., 2007). This approach has also been shown to have high scan-rescan reliability for fMRI (Schuyler et al., 2010).

We implemented DCM in SPM12. DCM presents an advantage over other measures of effective connectivity such as structural equation modeling (C. Büchel & Friston, 1997; McIntosh & Gonzalez-Lima, 1994) or Granger causality (Goebel et al., 2003) because it not only provides a measure of endogenous effective connectivity between regions, but also a measure of how these directed connections are modulated by task demands (Stephan & Friston, 2010). In DCM, the state equations are causal in form – meaning the rate of change in each region depends on the current state of the system (i.e., mechanistic effects; Chicarro & Ledberg, 2012). To be clear, by “mechanistic” here we mean plausible neuronal mechanisms that underlie experimental measurements of brain responses (Stephan et al., 2010). The resulting coupling parameters reflect effective connectivity: the estimated influence that one region exerts over another under different experimental conditions. When these parameters change across conditions, we interpret this as modulation of connectivity by external inputs or task conditions.

Experimental paradigms that maximize the discriminability between the different models are recommended for DCM (Daunizeau et al., 2011); our task design allowed us to make discrete comparisons between two factors, each with two levels: image type (i.e., scene vs. scrambled color-tile images) and viewing instructions (*free*-vs. *fixed*-viewing). The state equation for DCM is:

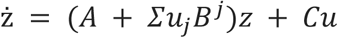

The intrinsic and baseline interactions between the neuronal states are endogenous connections and quantified by *A* parameters. These interactions are mediated by anatomical connections and are irrespective of task conditions (i.e., when the two conditions are combined). The influences of the task conditions on connectivity between ROIs are modulatory, and quantified by *B* parameters (i.e., connections altered on *free*-viewing encoding trials over-and-above all encoding trials). In this study, we used a mean-centered driving input so that the matrix *A* parameters represented an average effective connectivity across experimental conditions and matrix *B* modulatory parameters strengthen or weaken the average connectivity (Zeidman et al., 2019b). The influences of driving inputs are quantified by *C* parameters (in this case, caused by all encoding trials).

#### 2.8.1 Individual-level DCM analysis

Our main question of interest asked how unrestricted viewing during scene and scrambled image encoding modulated the effective connectivity architecture between the HPC, PPA, and OCP. Due to the increased complexity in interpretation when including inter-hemispheric connections for DCM (Stephan et al., 2010; Stephan & Friston, 2010; Zeidman et al., 2019b), we averaged across hemispheres for all models. Specifically, we configured the full model design with the HPC, PPA, and OCP to include all self-connections in the *A-*matrix (Figure 2A). Self-connections determine the excitatory/inhibitory balance within ROIs and provide a biologically-plausible mechanism for changes in the ROI’s activity through the interplay of interneurons and pyramidal cells (Bastos et al., 2012). We also included reciprocal connections between the PPA and OCP as well as between the PPA and HPC (Figure 2A) based on prior work suggesting likely structural connections (Shen et al., 2016). These self-connections and coupling parameters between ROIs were included in the modulatory *B*-matrix (Figure 2A), allowing us to examine our main research question, i.e., how unrestricted viewing modulated these connections. For simplicity, we set the input (*C*-matrix) to enter the earliest ROI in the visual stream (i.e., OCP). We conducted the first-level analysis by specifying a DCM of the full model design for each subject then estimating the model parameters, separately for the scene and scrambled encoding conditions. Finally, to confirm that potential lateralization differences in scene encoding (see e.g., Silson et al., 2019) would not be missed by averaging across hemispheres, we also modeled the *A*- and *B*-matrices in the left and right hemispheres separately for the scene encoding condition only.

**Figure 2.**
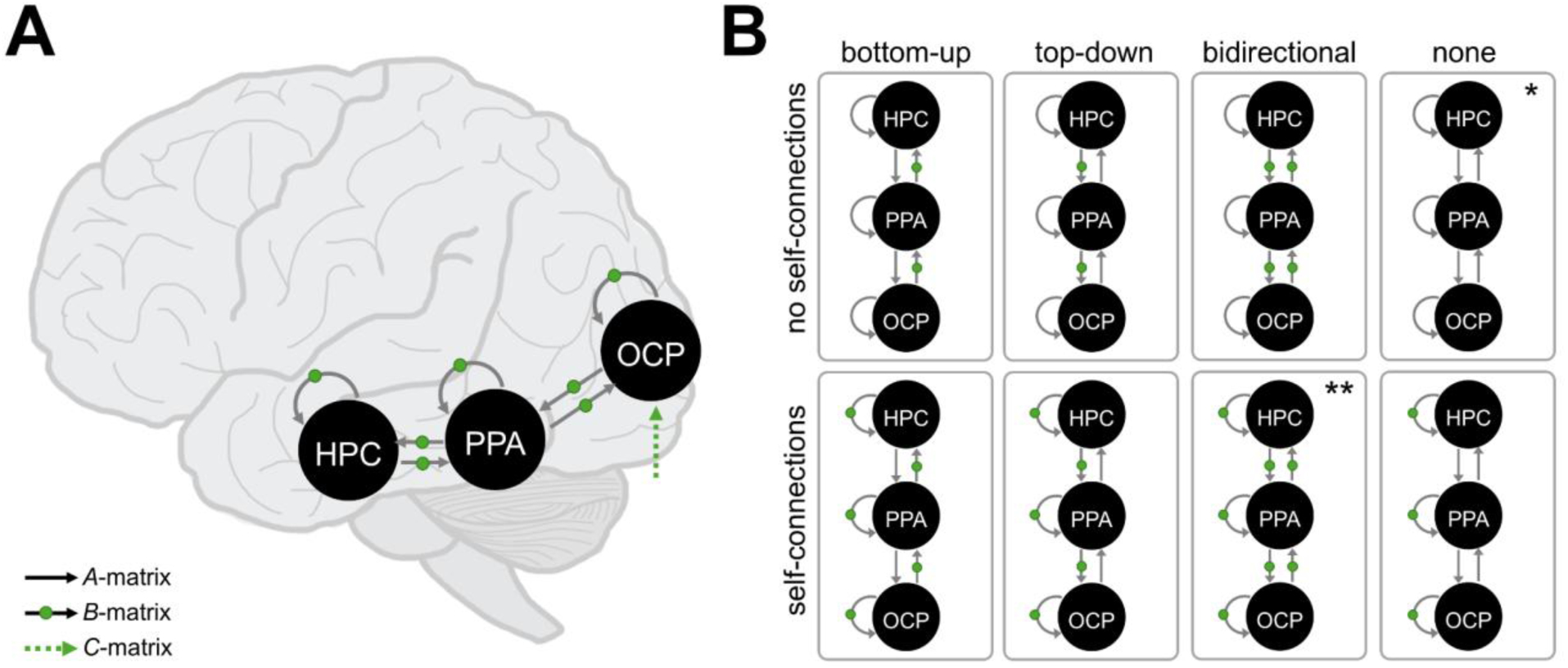
Model design for dynamic causal modeling (DCM) showing self- and coupling parameters for the hippocampus (HPC), parahippocampal place area (PPA) and occipital pole (OCP). **(A)** Full model estimated using Parametric Bayes (PEB) approach including endogenous (i.e., *A*-matrix) connections shown with black arrows, modulatory (i.e., *B*-matrix) connections shown with green dots, and input (i.e., *C*-matrix) shown with a dotted green line. **(B)** Model space consisting of 7 “families” based on the directionality of modulatory connections, plus the null model (indicated by *), evaluated using Bayesian model selection at the family-wise level. The full model family is indicated by **.

#### 2.8.2 Group-level DCM analysis

After specifying and estimating DCMs at the subject level, we used Parametric Empirical Bayes (PEB) analysis to evaluate group-level effects and between-subjects variability on parameters. PEB assumes that all subjects have the same basic architecture but differ in terms of the strength of connections within that model (Zeidman et al., 2019b).

For the *A*- and *C*-matrices, we then performed model comparison by running an automatic search with Bayesian Model Reduction (BMR). Here, parameters from the PEB model that do not contribute to model evidence are pruned (Penny et al., 2004, 2007). Finally, the parameters from the best reduced models were averaged using Bayesian Model Averaging (BMA) weighted by their evidence, which resulted in the winning models for scene and scrambled image encoding.

To test specific hypotheses about model parameters in the *B*-matrices, we first conducted Bayesian model comparison using a family-wise approach within a predefined model space (Penny et al., 2010; Figure 2B). Specifically, we investigated the effects of modulation by *free*-viewing for each of the two image encoding conditions by specifying a model space consisting of 7 “families” based on the directionality of connections: (1) bottom-up, (2) top-down, (3) bidirectional, (4) self-inhibitory, (5) bottom-up + self-inhibitory, (6) top-down + self-inhibitory, and (7) bidirectional + self-inhibitory.

The model space consisted of 128 models, including the full model (in which all self- and between-ROI connections were turned on) and the null model (in which all self- and between-ROI connections were turned off). These models were assessed for evidence using Bayesian model selection at the family-wise level, selecting the family of models with the greatest posterior probability (Pp). Parameters from the winning family of models were averaged using BMA, which resulted in the final winning models for *free*-viewing modulation of scene and scrambled image encoding.

In line with recommendations on reporting DCM results (see Stephan et al., 2010), here we focused on those parameters in the winning models with a posterior probability (Pp) equal to or greater than 95% (i.e., strong evidence; Kass & Raftery, 1995). Importantly, intrinsic (i.e., self-) connections are inhibitory by definition and are measured on a unitless log scale (Zeidman et al., 2019a) such that positive values reflect more self-inhibition and negative values reflect disinhibition (i.e., increased sensitivity to inputs). Coupling parameters (i.e., between-node connections) are measured in hertz, and so lend themselves to a simpler interpretation: if a connection is positive, then it is excitatory and if the connection is negative, then it is inhibitory.

To investigate the effect of meaningful content on modulatory effective connectivity strengths (Ep), we ran a 2×2 repeated measures ANOVA, specifying main effects and their interactions for image type (scene, scrambled) and DCM connection (OCP, PPA, HPC, OCP to PPA, HPC to PPA, PPA to OCP, PPA to HPC). The model was fit using the “aov_4” function in the *afex* package (Singmann et al., 2024) in R 4.4.2 (R Core Team, 2024). We used individual participants’ parameters following PEB procedures, which had been updated using the group-level connection strengths as empirical priors. This facilitated the primary analysis focused on planned pairwise comparisons between scene and scrambled image types within each connection. These comparisons of interest were examined using the “contrast” function in the *emmeans* package (Lenth, 2021). All *p*-values were adjusted by controlling for False Discovery Rate (FDR).

#### 2.8.3 Relationship with retrieval-related measures

In addition to assessing the group-level *B*-matrix modulatory results, we also investigated the relationship of the modulatory connections with two retrieval-related measures with each connection: (1) the average difference in subsequent memory retrieval strength between *free*- and *fixed*-viewing and (2) the average difference in gaze reinstatement similarity between *free*- and *fixed*-viewing. Here, we sought to answer how between-subject differences in recognition memory performance and memory-related gaze reinstatement for scenes could be explained by the differences in the excitability or inhibition of ROIs and their coupling parameters during encoding with unrestricted viewing.

After the fMRI encoding task (see 2.3 Procedure above), participants were given a 60-minute break before they began a retrieval task in a separate testing room. This post-scan retrieval task is described in detail in Liu et al. (2020) and is summarized here. All scene and scrambled images from the fMRI encoding task were tested in the retrieval task, along with 288 images that were not used in the encoding task as lures (total images = 864). These images were divided into 6 blocks. In each retrieval trial, a fixation cross “+” was presented for 1.5 seconds and this cross was presented at the same location as it was during the encoding task for previously viewed images. Then, the retrieval image was presented on the screen for 4 seconds. Participants were given 3 seconds to indicate whether the image was “old” or “new” (i.e., previously viewed during the fMRI task) using 4 keys on the keyboard: z (high confidence, old), x (low confidence, old), n (high confidence, new) and m (low confidence, new). During the retrieval task, participants’ eye movements were recorded using the Eyelink II head mounted infrared camera system (SR Research LTD., Ottawa, Ontario, Canada) at a sampling rate of 500 Hz.

Subsequent memory retrieval strength was calculated as follows: participants’ responses were assigned 2 points to stimuli that were correctly recognized as previously viewed with high confidence, 1 point to stimuli that were correctly recognized with low confidence, 0 points to previously viewed stimuli that were incorrectly endorsed as new with low confidence, and −1 point for previously viewed stimuli that were incorrectly endorsed as new with high confidence. As described in further detail in Wynn et al. (2022), the spatial overlap between gaze patterns elicited by the same participants viewing the same images during encoding and retrieval was quantified into gaze reinstatement scores. This was done by computing the Fisher *z*-transformed Pearson correlation between the duration-weighted fixation density map for each image for each participant during encoding and the corresponding density map for the same image viewed by the same participant during retrieval (R *eyesim* package: https://github.com/bbuchsbaum/eyesim; for further details regarding density map computation, see Wynn et al., 2020). To account for idiosyncratic viewing tendencies (e.g., participant’s tendency to view each image from left to right) we also computed the similarity between participant- and image-specific retrieval density maps and 50 other randomly selected encoding density maps from the same participant in the same condition, yielding an averaged single control score for each participant for each image. The difference between these two gaze similarity scores reflects gaze reinstatement similarity, i.e., spatial similarity between encoding and retrieval scanpaths for the same participant viewing the same image, controlling for idiosyncratic viewing biases.

For each participant, we calculated the difference in the average (i.e., across all trials) score for memory retrieval strength and gaze reinstatement similarity between *free*- and *fixed*-viewing of scenes, which were subsequently mean-centered.

We applied canonical variate analysis (CVA) to model the effect of the modulatory self- and between-region parameter strengths resulting from the DCM analysis against memory strength and gaze reinstatement similarity (i.e., retrieval-related measures) with unrestricted viewing for the scene condition only. In this way, we could assess whether greater differences in memory performance and gaze reinstatement between *free*- and *fixed*-viewing of scenes are associated with stronger modulatory effects during encoding with unrestricted viewing. CVA allowed dimension reduction of the parameters in the winning model for the *B*-matrix, so that we could investigate whether the behavioral measures were associated with the first derived canonical variate. We assessed canonical loadings (i.e., the correlation between each original variable and the canonical variates) to determine how variables contribute to each respective canonical variate while accounting for multicollinearity.

## 3 Results

For both scene and scrambled models, we found positive driving inputs (i.e., *C*-matrix) to the OCP (scene: 0.638, Pp ≥ 99%; scrambled: 0.173, Pp ≥ 99%).

### 3.1 Endogenous effective connectivity

We first established the winning models of endogenous (i.e., *A*-matrix) effective connectivity for scene and scrambled image encoding using a BMA approach, the results of which can be seen in Figures 3A and 3B, respectively. For scene encoding, we found positive, intrinsic (i.e., self-) connections for all ROIs (HPC: 0.207; PPA: 0.407; OCP: 0.218, *all* Pp ≥ 99%). All between-ROI connections were also favored, except for the connection from the PPA to the HPC (Pp = 28%). Specifically, BMA revealed negative top-down coupling parameters from the HPC to PPA (−0.277, Pp ≥ 99%) and from the PPA to OCP (−0.657, Pp ≥ 99%). Finally, we also found positive, bottom-up connectivity from the OCP to PPA (0.747, Pp ≥ 99%). Follow-up analyses in each hemisphere separately revealed no differences in the pattern of endogenous effective connectivity for scene encoding, except for the self-connection of the left HPC, which did not reach the 95% Pp threshold.

**Figure 3.**
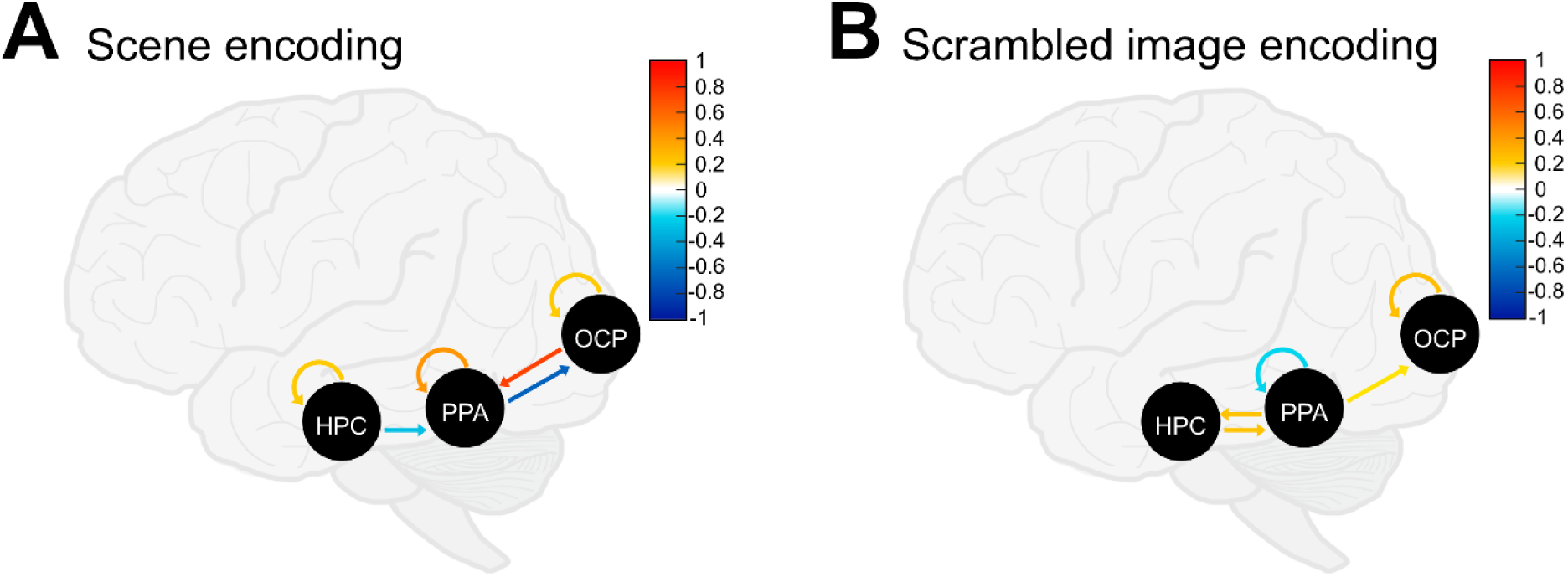
*A*-matrix (i.e., endogenous) effective connectivity results from Bayesian Model Averaging (BMA). (A) Scene image encoding model resulting from BMA. (B) Scrambled image encoding model resulting from BMA. For (A) and (B) self-connections shown in warm colors (i.e., yellow-red) are positive and reflect greater self-inhibition of the ROI, whereas self-connections shown in cool colors (i.e., blue) are negative and reflect greater disinhibition of the ROI. Between-node connections are shown in warm colors (i.e., yellow-red) for positive coupling parameters, and in cool colors (i.e., blue) for negative coupling parameters.

For scrambled image encoding, self-connections were found to be negative (i.e., self-disinhibitory) for the PPA (−0.179, Pp = 98%) and positive (i.e., self-inhibitory) for the OCP (0.264, Pp ≥ 99%). Note that the HPC self-connection failed to reach the 95% Pp threshold, but was nonetheless highly probable (−0.156, Pp = 93%). We found positive coupling parameters from the HPC to PPA (0.247, Pp ≥ 99%) and vice versa (0.235, Pp ≥ 99%). BMA did not result in bottom-up connectivity from the OCP to the PPA (0.000, Pp = 15%), instead finding positive top-down coupling from the PPA to OCP (0.125, Pp = 98%).

### 3.2 Modulatory connections

After establishing the winning models of endogenous connectivity for scene and scrambled image encoding, we next assessed family-level evidence for modulatory (i.e., *B*-matrix) effects of *free*-viewing. The winning model family for scene encoding modulated by *free*-viewing was the family that included self-connections and bidirectional between-ROI connections (Pp ≥ 99%, indicated by ** in Figure 2B). We found little or no evidence for any other model family for modulatory effects of *free*-viewing on scene encoding. Likewise, the winning model family for scrambled image encoding modulated by *free*-viewing also included bidirectional couplings and self-connections (Pp ≥ 99%). We found weak evidence for scrambled image encoding modulated by *free*-viewing of the top-down + self-inhibitory model family (Pp = 7%). This is considered low evidence, so this family was not considered in subsequent BMA, averaging only within the winning model family.

The averaged best candidate models within the winning family for the modulatory effect of *free*-viewing on scene encoding included all self-connections and all between-ROI connections, except for the connection from the PPA to the OCP (Figure 4A). Specifically, we found negative, i.e., self-excitatory, modulation by *free*-viewing on the OCP (−1.277, Pp ≥ 99%), the PPA (−0.744, Pp ≥ 99%) and the HPC (−0.867, Pp ≥ 99%). Turning to the coupling parameters, we found a top-down, inhibitory effect of *free*-viewing on scene encoding from the HPC to the PPA (−2.44, Pp ≥ 99%), which did not extend from the PPA to the OCP (Pp = 21%). Additionally, there was a bottom-up, positive modulatory effect (i.e., excitatory) of *free*-viewing extending from the OCP to the PPA (0.346, Pp ≥ 99%), and from the PPA to the HPC (0.262, Pp ≥ 99%) for scene encoding. Follow-up analyses in each hemisphere separately revealed no differences in the pattern of modulatory effective connectivity for scene encoding.

**Figure 4.**
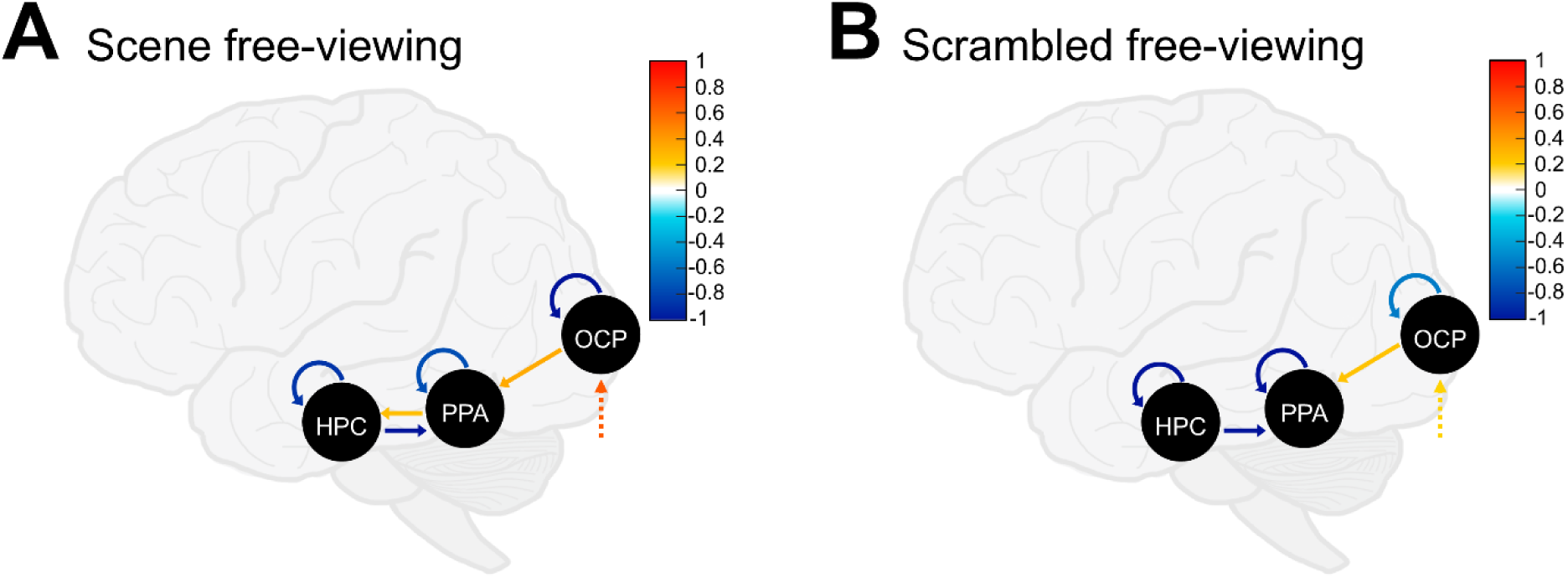
*B*-matrix (i.e., modulatory effects) effective connectivity results from Bayesian Model Averaging (BMA). Input (*C*-matrix) is shown with dashed arrows to the occipital pole (OCP) in both models. (A) Scene image encoding modulated by *free*-viewing resulting from BMA. (B) Scrambled image encoding modulated by *free*-viewing resulting from BMA. For (A) and (B) self-connections shown in warm colors (i.e., yellow-red) are positive and reflect greater self-inhibition of the ROI, whereas self-connections shown in cool colors (i.e., blue) are negative and reflect greater disinhibition of the ROI. Between-node connections and inputs are shown in warm colors (i.e., yellow-red) for positive coupling parameters, and in cool colors (i.e., blue) for negative coupling parameters.

For scrambled image encoding, the averaged best candidate models within the winning family for the modulatory effect of *free*-viewing included all self-connections and two between-ROI connections (Figure 4B). Specifically, we found negative (self-excitatory) modulation by *free*-viewing on the OCP (−0.539, Pp ≥ 99%), the PPA (−2.003, Pp ≥ 99%), and the HPC (−1.102, Pp ≥ 99%). Only two coupling parameters (i.e., between-ROI connections) survived: a top-down, inhibitory effect of *free*-viewing on scrambled image encoding from the HPC to the PPA (−1.100, Pp ≥ 99%), and a bottom-up, excitatory modulatory effect from the OCP to the PPA (0.241, Pp > 95%).

A 2×2 repeated measures ANOVA (image type by DCM connection) on modulatory effective connectivity strength (Ep) found a main effect of image type (*F*_2.23,78.10_ = 223.73, *p* < .001), which was qualified by an interaction with DCM connection (*F*_2.13,74.67_ = 57.79, *p* < .001). Estimated marginal means were calculated for the effect of image type across each level of DCM connection, summarized in Table 2. For self-connections, Ep modulated by *free*-viewing was significantly more negative for scenes compared to scrambled images in the OCP (*p* < .001), reflecting greater self-disinhibition for scene images. In contrast, Ep modulated by *free*-viewing was significantly less negative for scenes compared to scrambled images in the PPA (*p* < .001) and in the HPC (*p* < .01), reflecting greater self-inhibition for scene images. For the coupling parameter from the HPC to the PPA, Ep modulated by *free*-viewing was significantly more negative (i.e., inhibitory) for scenes compared to scrambled images (*p* < .001). Finally, Ep was significantly more positively modulated by *free*-viewing in the coupling parameter from the PPA to the HPC for scenes compared to scrambled images (*p* < .05). No other comparisons were significant (all *p*s > .05).

**Table 2.** Differences in estimated marginal means for the effect of image type by DCM connection. (*indicates *p < .05*).

| **Contrast**<br>**(Scene-Scrambled)** | **Estimate** | **t-ratio** | **p** |
| --- | --- | --- | --- |
| OCP to OCP | -0.728 | --4.893 | <.001* |
| PPA to PPA | 1.266 | 18.108 | <.001* |
| HPC to HPC | 0.162 | 3.423 | .003* |
| OCP to PPA | 0.105 | 1.078 | .337 |
| PPA to OCP | -0.021 | -0.120 | .905 |
| PPA to HPC | 0.128 | 2.712 | .014* |
| HPC to PPA | -1.3855 | -12.154 | <.001* |

### 3.3 Relationship with memory strength and gaze reinstatement

In addition to the average modulatory effect of *free*-viewing on encoding, we were interested in quantifying the relationship of two retrieval-related measures (average difference in subsequent memory strength and gaze reinstatement similarity for scenes encoded in *free-* over *fixed-*viewing conditions) with each DCM modulatory connectivity parameter. This analysis included modulatory parameters that reached Pp ≥ 95% in BMA. We found that the canonical variate derived from the DCM connectivity parameters correlated significantly with the canonical variate derived from the retrieval-related measures [*r* = 0.62 (Pearson’s correlation coefficient), *p* < .001], indicating that 38.4% of the shared variance was explained by the first canonical function. Canonical loadings revealed that greater self-connectivity of the PPA (*r* = 0.67) and HPC (*r* = 0.80) were the strongest contributors to the canonical variate, explaining 44.7% and 64.3% of the variance, respectively. This was associated primarily with subsequent memory strength (*r* = 0.89) but the canonical loading also revealed a moderate contribution of gaze reinstatement similarity (*r* = 0.34). In other words, greater self-connectivity (i.e., less disinhibition, or less receptivity to input from the model) of the PPA and HPC was significantly associated with greater subsequent memory strength and greater gaze reinstatement similarity for *free*-compared to *fixed*-viewing of scenes. Importantly, even when the endogenous (i.e., *A*-matrix) connectivity parameters that reached 95% Pp were added to the model, greater modulatory self-connectivity of the PPA and HPC continued to be the strongest contributors to the canonical variate – but not their endogenous self-connections.

## 4 Discussion

Memory and gaze behavior are thought to be intricately linked (Ryan & Shen, 2020; Yarbus, 1967), guiding one another in service of ongoing encoding and retrieval processes. Thus, the neural architecture of the brain must support rapid, reciprocal exchange of information between the extended HPC memory and visuo-oculomotor control systems. Modeling of directed connections in the macaque brain has revealed likely anatomical pathways from the HPC to visuo-oculomotor systems (Shen et al., 2016) and also how these connections are functionally relevant to facilitating the flow of information from the HPC about mnemonic representations to influence visuo-oculomotor regions (Ryan et al., 2020). Convergent evidence from human neuroimaging has demonstrated that these systems work together in service of building mental representations of scenes for later retrieval (Liu et al., 2020) and that information about patterns of gaze behavior at encoding may be embedded in these mental representations to be recapitulated at retrieval (Wynn et al., 2022). Critically, however, evidence for directionality of information flow in humans during encoding within this reciprocal architecture is so far limited because measures of functional connectivity can only reveal the degree to which component regions correlate over time. The current study uses a measure of effective connectivity to provide a non-invasive account for the directional influences between these systems when new memories are created, and— for the first time—how unrestricted, naturalistic visual exploration induces changes in this connectivity. Importantly, the current findings also suggest that this pattern of effective connectivity modulated by unrestricted viewing predicts subsequent memory retrieval performance, demonstrating a biologically-plausible model for the mechanism underlying a rich history of findings starting with seminal work by Loftus (1972) linking greater gaze fixations to better subsequent memory (Henderson et al., 2005; Liu et al., 2017; Olsen et al., 2016). Altogether, these findings highlight how gaze behavior shapes the ways in which brain dynamics within and between the extended HPC memory system and visuo-oculomotor control system unfold during encoding in humans, extending earlier neuroimaging findings focused on functional measures of connectivity (Liu et al., 2020; Wynn et al., 2022).

The PPA is well-connected with regions of the visuo-oculomotor system to receive and process incoming visuo-spatial information (Baldassano et al., 2013) and may reflect maintained retinotopic organization along the visual stream (Huang & Sereno, 2013; Silson et al., 2015). Importantly, prior functional connectivity analyses of the current dataset revealed that unrestricted visual exploration – compared to when viewing is restricted – enhances functional activation during scene encoding along the ventral visual stream and into the PPA (Liu et al., 2020). Extending these findings, here we show how excitatory effective (i.e., directional) connectivity from early visual cortex to the PPA is enhanced by unrestricted visual exploration. Specifically, during scene encoding, we found bottom-up, excitatory endogenous connectivity from the OCP to the PPA and top-down, inhibitory connectivity from the PPA to the OCP across viewing conditions. In line with our predictions, *free*-viewing further enhanced this bottom-up connectivity and also disinhibited self-connections in both the PPA and OCP, thus increasing the sensitivity of both regions to input coming from the rest of the network (Bastos et al., 2012). Indeed, the OCP has been implicated in sampling informative regions at encoding (Fehlmann et al., 2020) and in the formation of gaze scanpaths (Wynn et al., 2022), suggesting that it may play a role in synchronizing visual information across different gaze locations which are then integrated into a coherent mental representation of space at the level of the PPA (Golomb et al., 2011). Increased excitatory connectivity from early visual cortex to the PPA during *free*-viewing is also consistent with an interpretation under predictive coding models (e.g., de-Wit et al., 2010) which suggests that the brain actively anticipates upcoming sensory input rather than passively registering it. Although we did not explicitly manipulate or measure predictive errors in the current study, there is mounting evidence that the process of prediction enhances encoding (Poskanzer et al., 2025), and unrestricted viewing in the *free*-viewing condition may have allowed participants to test predictions about what features should be present within the scenes. In contrast, the inhibitory endogenous connection from PPA to OCP during scene encoding may represent a suppression of top-down signal, reflecting a balancing mechanism to instead facilitate the positive, bottom-up signal towards the extended HPC memory system (Zeidman et al., 2019b). Alternatively, it could be argued that the pattern of effective connectivity between the OCP and PPA simply reflects relative differences in attentional or working memory demands induced by task instructions. However, prior work suggests that maintaining central fixation (i.e., *fixed*-viewing) does not significantly increase these demands (Armson et al., 2019), perhaps because participants still move their eyes, albeit with fewer fixations and smaller saccade amplitudes, consistent with other work in the literature (Damiano & Walther, 2019; Liu et al., 2020; Welke & Vessel, 2022). Moreover, in examining the total viewing time that was directed to the scenes, we found that participants spent as much, if not more, total viewing time looking at the scenes in the *fixed*-viewing condition compared to the *free*-viewing condition. If participants had differentially disengaged their attention from the *fixed*-viewing condition, we may have expected to see more off-screen viewing behavior, resulting in a lower total duration of viewing to the scenes. Instead, attentional engagement, operationalized as looking time, remained comparable during fixed- and *free*-viewing, but the amount of information extracted through gaze fixations differed.

Within the extended HPC memory system, *free*-viewing during scene encoding increases connectivity between the PPA and HPC (Liu et al., 2020). Specifically, in line with our hypotheses, we found that bottom-up modulatory excitatory connectivity driven by input to the OCP extended to the PPA and ultimately to the HPC, facilitating the flow of information accumulated during visual exploration into mental representations of scenes. Neural states of the PPA have recently been tied to both low-level visual features and large-scale spatial locations (Oetringer et al., 2025), making it a prime candidate region for the transfer and processing of visual input towards coherent mental representations. Our results further characterized the modulatory effect of unrestricted viewing on this connective architecture by also identifying a disinhibitory response within the HPC and an inhibitory connection from the HPC to the PPA. One possibility is that increased disinhibition of the HPC, in response to excitatory input from the PPA, may reflect a novelty response to scenes that thereby signals to encode, rather than to strengthen, an existing representation. A similar result was reported by Schott et al. (2023), in which the authors reported self-disinhibition of the HPC in response to input from the PPA during novel scene encoding, and here we show that this input is carried from the early visual cortex in response to unrestricted gaze behavior. At the cellular level, there is evidence that disinhibitory interneurons in specific HPC subfields may contribute to spatial learning (e.g., Turi et al., 2019). Nonetheless, the role of inhibitory/disinhibitory dynamics in the HPC during complex cognitive processes, such as memory encoding, is still a nascent investigation (Tzilivaki et al., 2023). Importantly, self-disinhibition of the HPC and PPA modulated by *free*-viewing was actually weaker for scenes compared to scrambled images suggesting that in order to encode and organize meaningful content in mind, some degree of self-inhibition in these regions is necessary. In other words, relatively less self-disinhibition for scenes may reflect higher levels of internal processing, facilitating a transition from active learning to consolidation processes of existing representations in memory (Büchel et al., 2024). Furthermore, we found inhibitory connectivity during scene encoding modulated by *free*-viewing from the HPC to the PPA, which may support a pattern separation-like mechanism to distinguish between overlapping features of previously viewed scenes within the task (Kirwan & Stark, 2007; Yassa & Stark, 2011). This interpretation could be further interrogated in a future study by explicitly manipulating the visual similarity of trial-adjacent scenes. We would predict greater inhibition of HPC to PPA signal when highly similar scenes are presented together, thereby necessitating greater separation of overlapping features in memory. Altogether, the winning model of scene encoding modulated by *free*-viewing is consistent with the notion that visual exploration supports increased information exchange between the visuo-oculomotor and extended HPC systems (Shen et al., 2016; Ryan et al., 2020).

Although we found some similarities in how unrestricted viewing modulated encoding of scenes and scrambled, color-tile images, there were two notable differences which warrant closer investigation: (1) endogenous effective connectivity across viewing conditions for scrambled image encoding was characterized by an excitatory connection from the PPA to the OCP; and (2) modulatory effective connectivity by unrestricted viewing did not extend to the HPC for scrambled image encoding. One potential interpretation of the former finding is that the PPA sends top-down signals to early visual cortex during scrambled image encoding based on prior knowledge and expectations about the content of scenes. Indeed, prior work has demonstrated that the PPA maintains functional responses to scrambled images of scenes (Coggan et al., 2019; Watson et al., 2017), confirmed in analyses of univariate activation with the current dataset (Liu et al., 2020). Additionally, in the absence of meaningful visual input, we previously identified memory-related top-down, excitatory effective connectivity from the PPA toward early visual cortex during scene construction (Ladyka-Wojcik et al., 2021). The latter finding lends itself to a more straightforward explanation: excitatory signals related to unrestricted viewing from the OPC towards the PPA did not extend to the HPC for scrambled image encoding because there was insufficient relevant visuo-spatial information to be extracted and bound into mental representations for later retrieval (Olsen et al., 2012; Yonelinas, 2013). Importantly, this key difference between scene and scrambled image encoding in the modulatory effect of *free*-viewing speaks to questions regarding the type of information that is carried via gaze fixations and on which the HPC operates (Hannula et al., 2010; Ryan, 2024; Voss et al., 2017). Effective connectivity during encoding may only propagate from the visuo-oculomotor system to the HPC when gaze behavior extracts meaningful information, such as the spatial and temporal arrangements of scene features (e.g., Vericel et al., 2024). More broadly, the visuo-oculomotor system seems to prioritize meaningful scene regions over bottom-up visual salience, even during tasks for which meaning is irrelevant for behavioral performance (Peacock et al., 2019). Since the scrambled images were matched to scene images in terms of low-level features, but lacked coherent spatial structure, it is likely that the OCP prioritized scene-relevant information that propagated through the connective architecture from the PPA to the HPC for successful encoding. Looking beyond the PPA, future studies could examine how effective connectivity of other nodes within the extended HPC system is dynamically modulated by viewing behavior, particularly in MTL regions, such as the entorhinal cortex, that are involved in spatial processing (Fyhn et al., 2004; Killian et al., 2012; Witter & Moser, 2006).

Canonical variate analysis revealed a strong relationship in our data between retrieval-related measures (i.e., memory strength and gaze reinstatement) and the effective modulatory parameter resulting from DCM for the *free*-viewing of scenes. The finding of such a relationship dovetails well with previous findings reported by Wynn et al. (2022) in which overlapping subsequent memory and gaze reinstatement modulation effects were found in the HPC memory system and in early visual cortex via cross-voxel brain pattern similarity analysis. Importantly, this relationship suggests that eye movement scanpaths at encoding not only facilitate the intake of visual information from our environment, but also are themselves built into memory representations at the level of the HPC system such that they can be leveraged to help with later retrieval. A closer inspection of canonical loadings revealed strong positive contributions of modulatory self-connections within the HPC and PPA (i.e., less disinhibition) – but not their endogenous self-connections – towards predicting a greater difference in retrieval-related performance with unrestricted viewing compared to restricted viewing. At the cellular level, inhibition during encoding may prevent excessive, runaway outbursts of postsynaptic activity (Kahan & Foltynie, 2013) and promote the consolidation and storage of memories for later retrieval (Barron et al., 2017). For example, GABAergic inhibitory interneurons in the HPC are thought to suppress irrelevant incoming activity to allow for more refined mnemonic representations (Pelkey et al., 2017) and play a fundamental role in further organizing these representations (McKenzie, 2018). In the current study, it may be the case that participants who had less modulatory self-disinhibition (i.e., greater self-inhibition) in these regions also showed greater differences in memory and gaze reinstatement for *free*-over *fixed*-viewing because they suppressed additional, irrelevant incoming visual information as they rapidly encoded and organized mnemonic representations of meaningful information (Katz et al., 2022).

On the other hand, participants with greater self-disinhibition of the HPC and PPA in response to unrestricted viewing of scenes may show less of a representational shift from encoding incoming visual information about scenes towards consolidation (Büchel et al., 2024). In rodents, co-occurring signals in the HPC during periods of rest associated with replaying past experiences and simulating future actions are thought to help the animal transition towards memory-guided action (Diba & Buzsáki, 2007; Pastalkova et al., 2008) and activity spikes within the HPC may even represent a demarcation of event boundaries between contexts during learning (Bulkin et al., 2020). At a systems-level, however, our understanding as to how the interplay of excitatory and inhibitory activation in the HPC enables encoding and later retrieval still needs greater investigation (Takeuchi et al., 2014). Future work should look to emerging evidence in the rodent literature examining HPC inhibition using encoding paradigms (Topolnik & Tamboli, 2022).

One limitation inherent to DCM approaches for fMRI data is the need to average within- and across-trials due to low temporal resolution, thus restricting the number of data points used to estimate parameters (David et al., 2006, 2008). As such, future work to disentangle the relationship between self- and coupling parameters at encoding with subsequent memory retrieval may benefit from innovative trial-by-trial approaches to DCM, using event-related potentials (see e.g., Asadpour & Wong-Lin, 2025). Trial-by-trial approaches paired with DCM for magnetoencephalography (Kiebel et al., 2009) may be particularly well-suited to our experimental task design, as the relationship between neural dynamics and gaze fixations likely fluctuates within the period of a single trial (Katz et al., 2020).

Furthermore, it should be noted that because DCM analyses are restricted to *a priori* hypothesized ROIs (Stephan et al., 2010), there may be additional higher-level cortical regions contributing to the excitatory-inhibitory balance within the extended HPC memory system that explain the relationship with retrieval-related measures found here. In a similar vein, future investigations could also probe the dynamics of connectivity parameters within subregions of the HPC (Hainmueller et al., 2024; McKenzie, 2018) and along the posterior-anterior axis of the PPA (Baldassano et al., 2013; Watson & Andrews, 2024), which may be associated with different aspects of subsequent mnemonic retrieval performance.

## 5 Conclusions

The neural architecture of the human brain must support the reciprocal exchange of information—including just-extracted information from visual exploration and previously stored mental representations—during encoding. In this study, we identified a biologically-plausible pathway in humans, dynamically modulated by gaze behavior, along which scene-relevant features can travel between the early visual cortex and the MTL. Active viewing enhances connections between these systems and predicts subsequent memory and replay of gaze patterns at retrieval. More broadly, our results demonstrate that the extended HPC memory and visuo-oculomotor control systems are intrinsically linked, and their directional interactions support the building of coherent, lasting mental representations.

## Data and Code Availability

Analysis scripts and final results matrices are openly available on Open Science Framework (OSF) at https://doi.org/10.17605/OSF.IO/SEUQN. All processed data associated with reported results are available on request from the corresponding author (NLW). The original data is not publicly available due to ethics restrictions at the time of data collection.

## Author Contributions

NLW: Conceptualization; Data curation; Formal analysis; Investigation; Methodology; Validation; Visualization; Writing – Original draft; Writing – Review.& editing. ZXL: Conceptualization; Data curation; Formal analysis; Investigation; Supervision; Methodology; Validation; Visualization; Writing – Original draft, Writing – Review & editing. JDR: Conceptualization; Funding acquisition, Project administration; Resources; Supervision; Writing – Original Draft; Writing – Review & editing.

## Acknowledgements

We would like to thank Arber Kacollja, Sarah Berger, Ling Li, Mandy Ding, Veena Sanmugananthan, Junghyun Nam, Mariam Aziz, and Jordana Wynn for their help at different stages of this research project.

